## Supplementary Information for "Comparing retinotopic maps of children and adults reveals a late-stage change in how V1 samples the visual field"

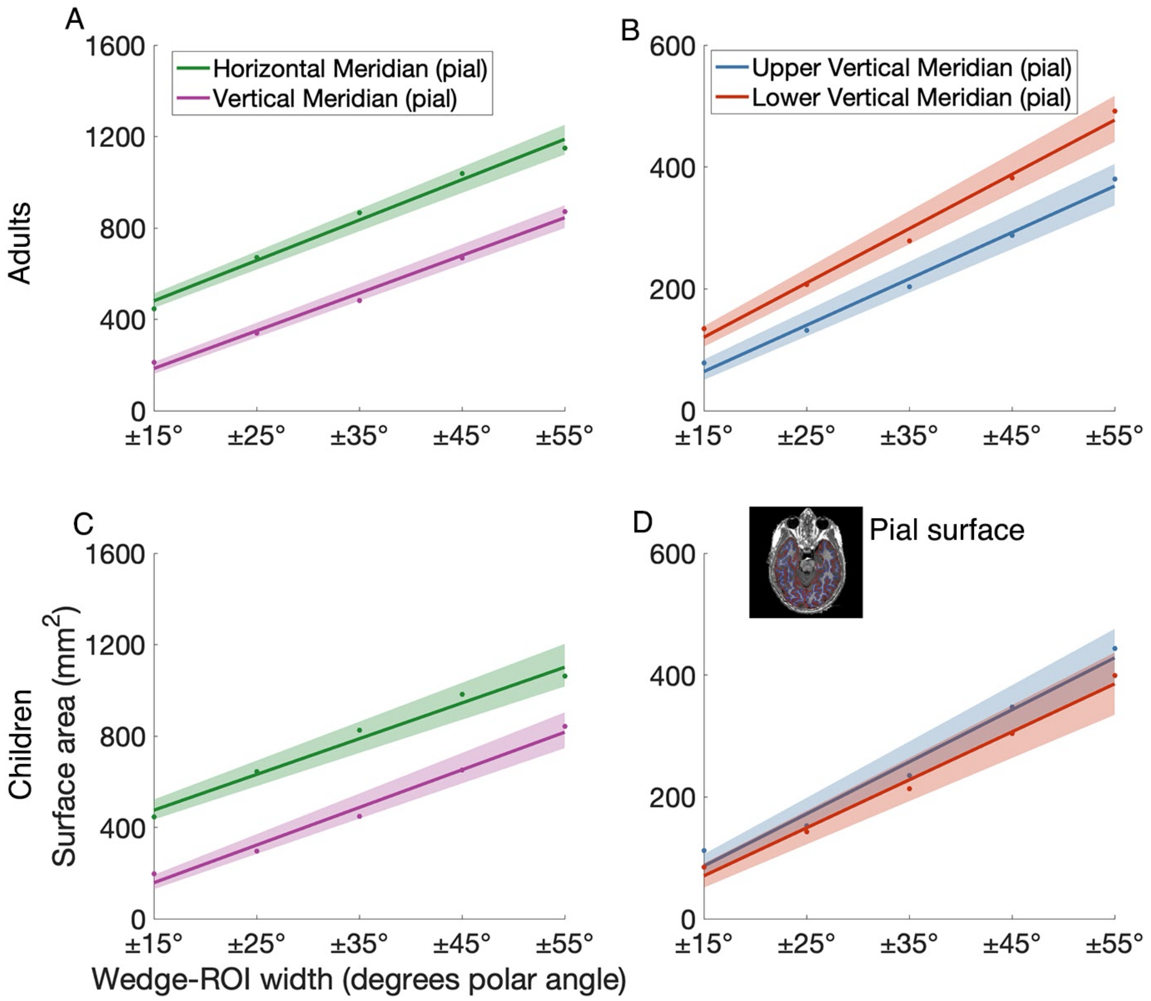


**SUPPLEMENTARY FIG. 1. (A-B)** V1 pial surface area (mm$^2$) measurements for wedges centered on the horizontal and vertical meridians, plotted as a function of wedge-width, for adults (n=24) and children (n=25). **(C-D)** V1 pial surface area (mm$^2$) measurements for wedges centered on the upper and lower vertical meridians, plotted as a function of wedge-width, for adults (n=24) and children (n=25). **(A-D)** Each line represents the average of 1000 bootstrapped linear fits to the data. The shaded error bar represents the 68% bootstrapped CI of the linear fit to the data. The inset in panel **(D)** illustrates an example of T1 anatomy scan with the white matter surface traced in blue and the pial surface traced in red.

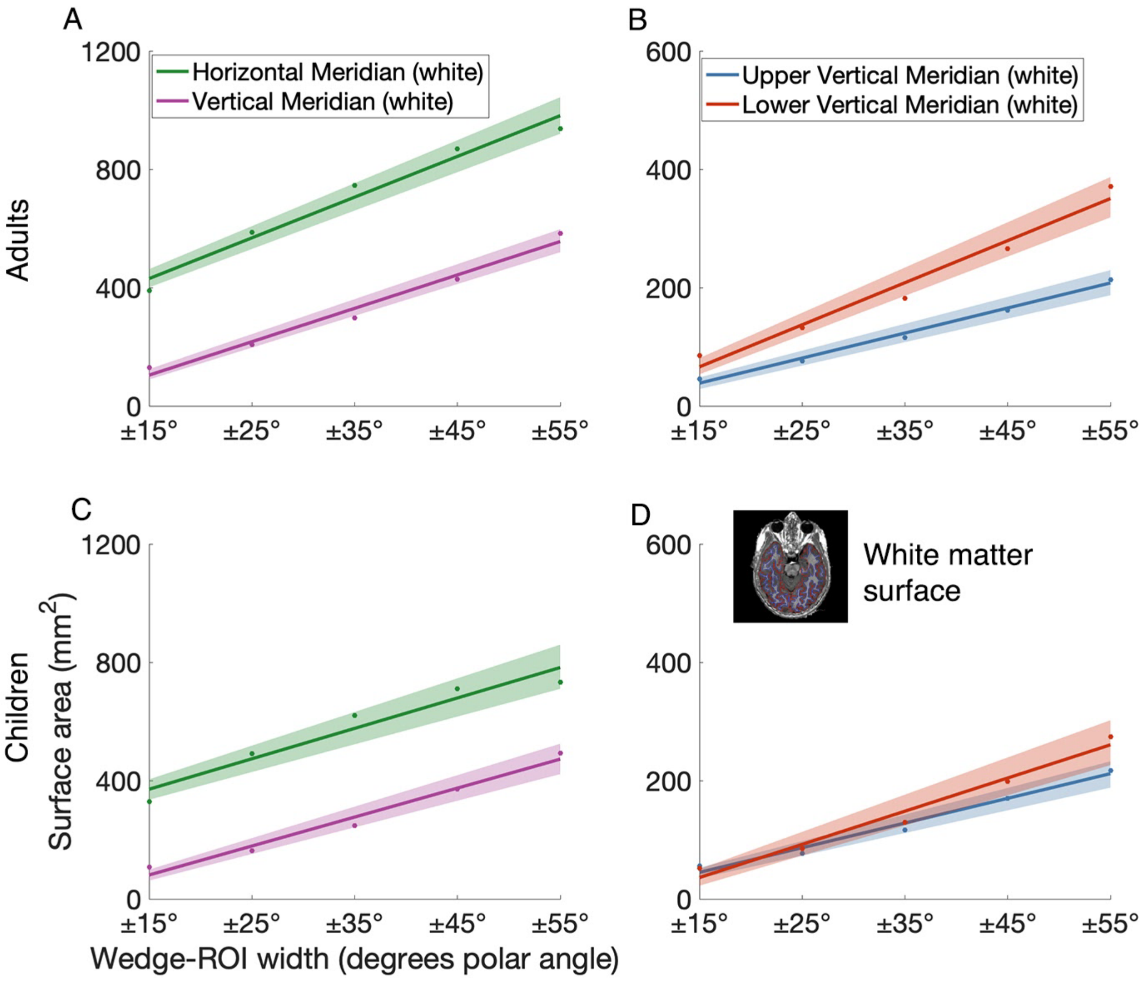


**SUPPLEMENTARY FIG. 2. (A-B)** V1 white matter surface area (mm$^2$) measurements for wedges centered on the horizontal and vertical meridians, plotted as a function of wedge-width, for adults (n=24) and children (n=25). **(C-D)** V1 white matter surface area (mm$^2$) measurements for wedges centered on the upper and lower vertical meridians, plotted as a function of wedge-width, for adults (n=24) and children (n=25). **(A-D)** Each line represents the average of 1000 bootstrapped linear fits to the data. The shaded error bar represents the 68% bootstrapped CI of the linear fit to the data. The inset in panel **(D)** illustrates an example of T1 anatomy scan with the white matter surface traced in blue, and the pial surface traced in red.

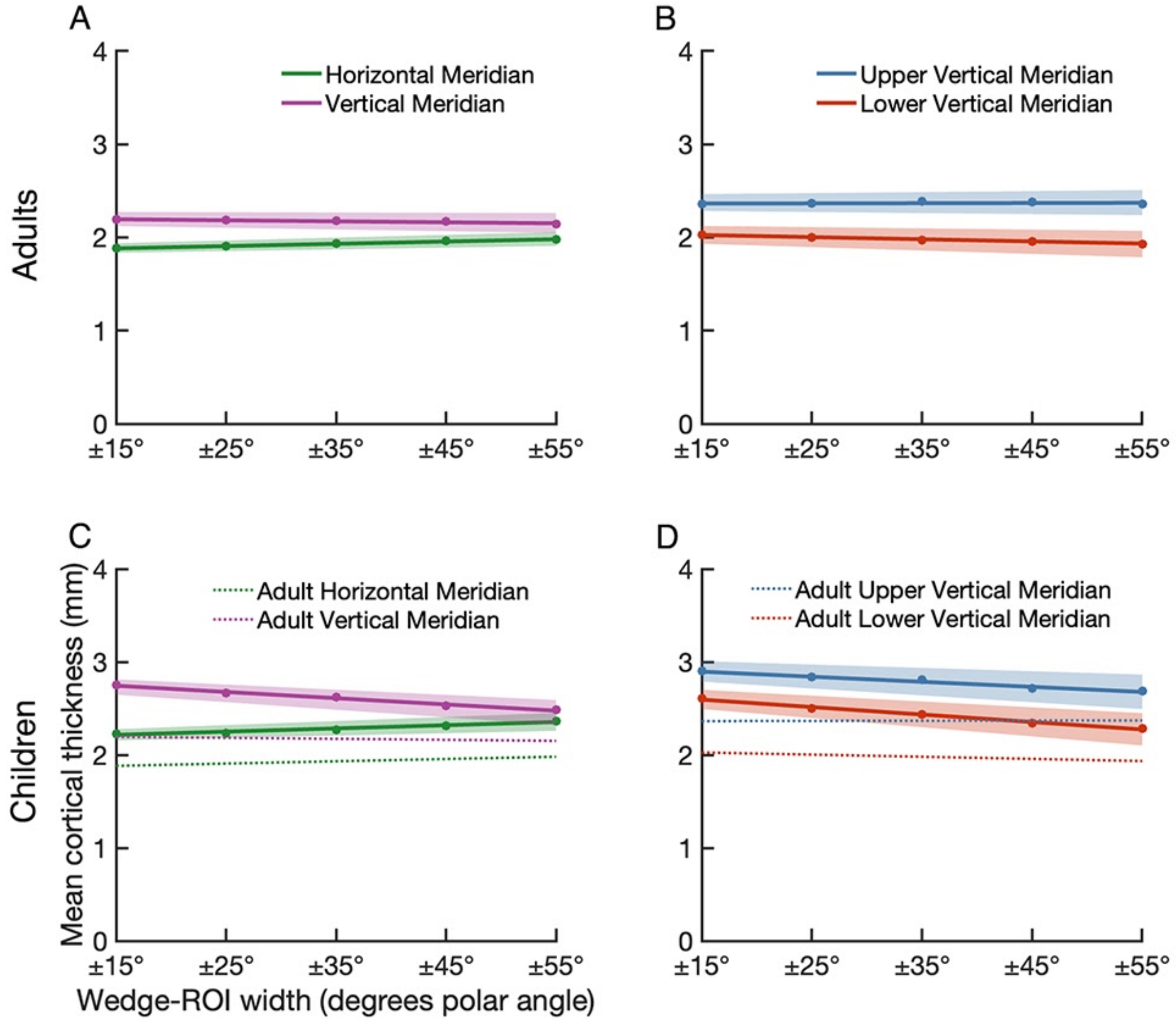


**SUPPLEMENTARY FIG. 3. (A-B)** V1 mean midgray thickness (mm) measurements for wedges centered on the horizontal and vertical meridians, plotted as a function of wedge-width, for adults (n=24) and children (n=25). **(C-D)** V1 mean midgray thickness (mm) measurements for wedges centered on the upper and lower vertical meridians, plotted as a function of wedge-width, for adults (n=24) and children (n=25). Each line represents the average of 1000 bootstrapped linear fits to the data. The shaded error bar represents the 68% bootstrapped CI of the linear fit to the data.

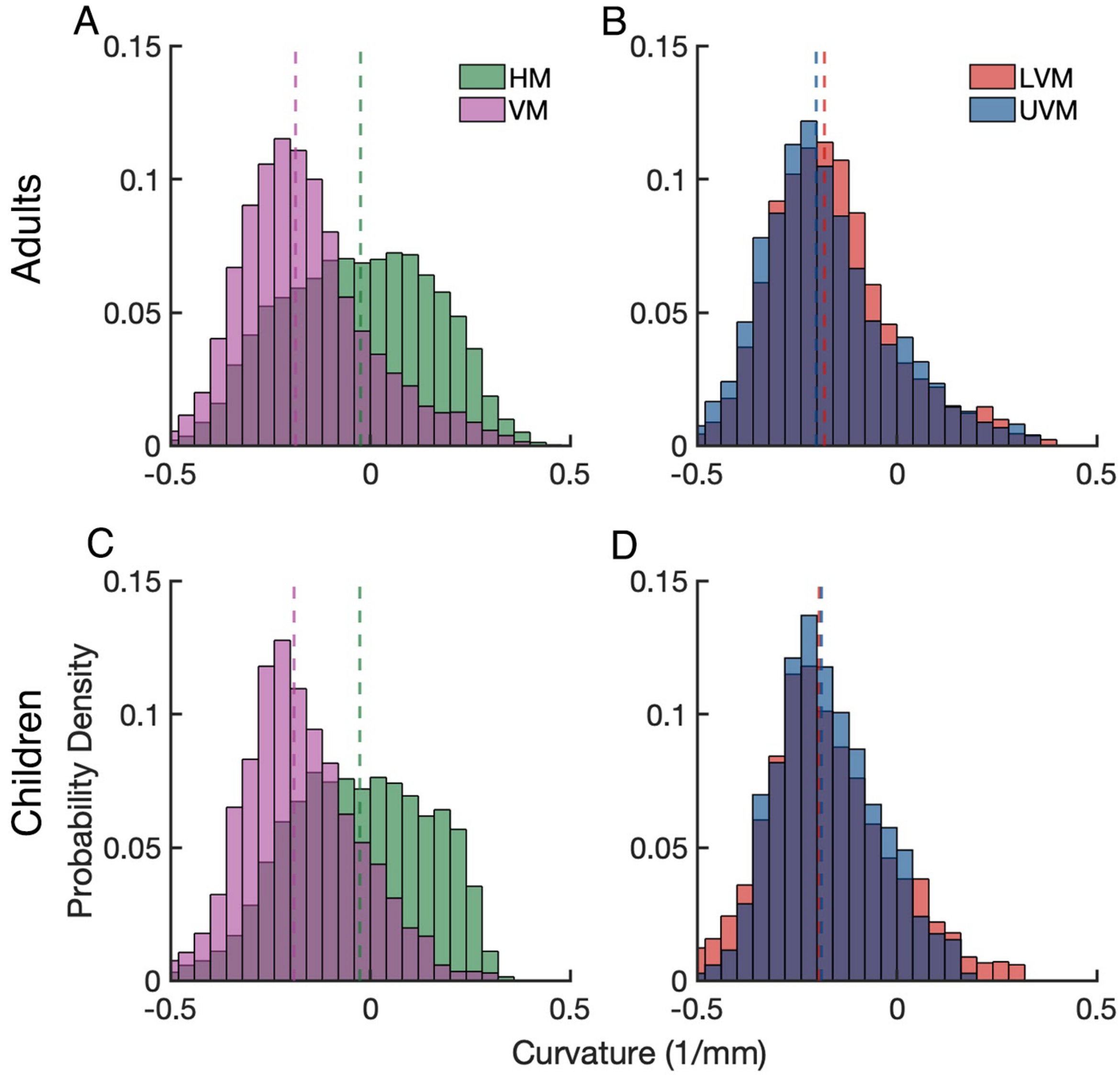


**SUPPLEMENTARY FIG 4.** Probability density histograms of vertex curvature values for ±25° wedges centered on the horizontal and vertical meridians, plotted as a function of wedge-width, for adults (n=24) and children (n=25). Dashed colored lines indicate the median curvature for each meridian. For adults, the median curvature for the horizontal meridian was -0.02, and for the vertical meridian it was -0.19. The lower vertical meridian had a median curvature of -0.18 and for the upper vertical meridian it was -0.20. For children, the horizontal meridian had a median curvature of -0.3 and for the vertical meridian it was -0.19. The lower vertical meridian had a median curvature of -0.19 and for the upper vertical meridian it was -0.19.

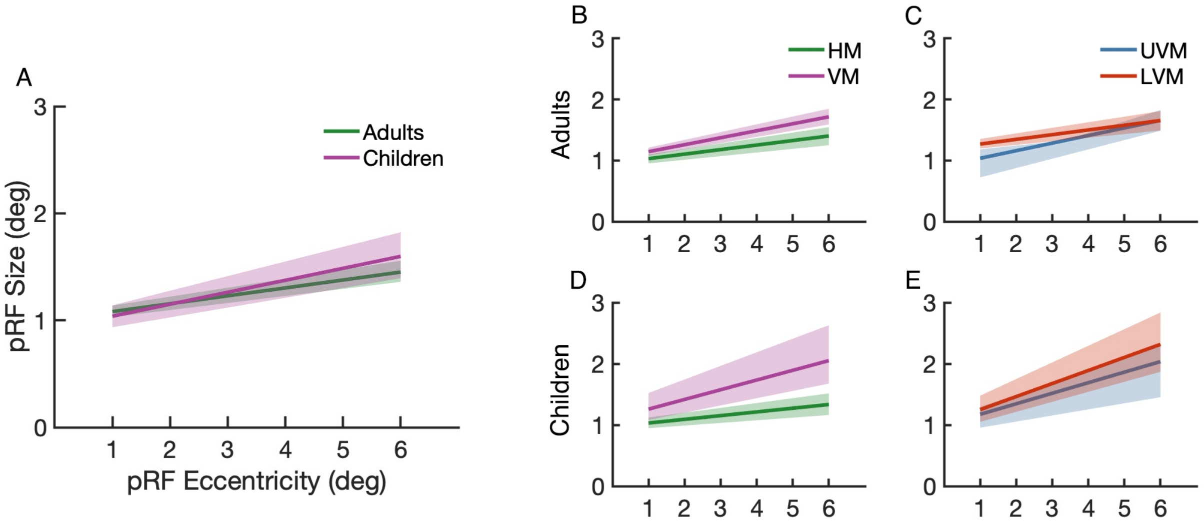


**SUPPLEMENTARY FIG 5**. Comparisons of median V1 pRF size between meridians for adults (n=24) and children (n=25). pRF size is plotted as a function of 1-6° eccentricity. **(A)** Comparisons of pRF size for adults and children across all of V1; **(B, D)** Comparisons of pRF size between the horizontal and vertical meridian for adults and children; **(C-E)** Comparisons of pRF size between the upper and lower vertical meridian for adults and children. Colored lines represent the average of 10,000 bootstrapped linear fits to the data. The shaded error bar represents the 68% bootstrapped confidence interval of a linear fit to the data.

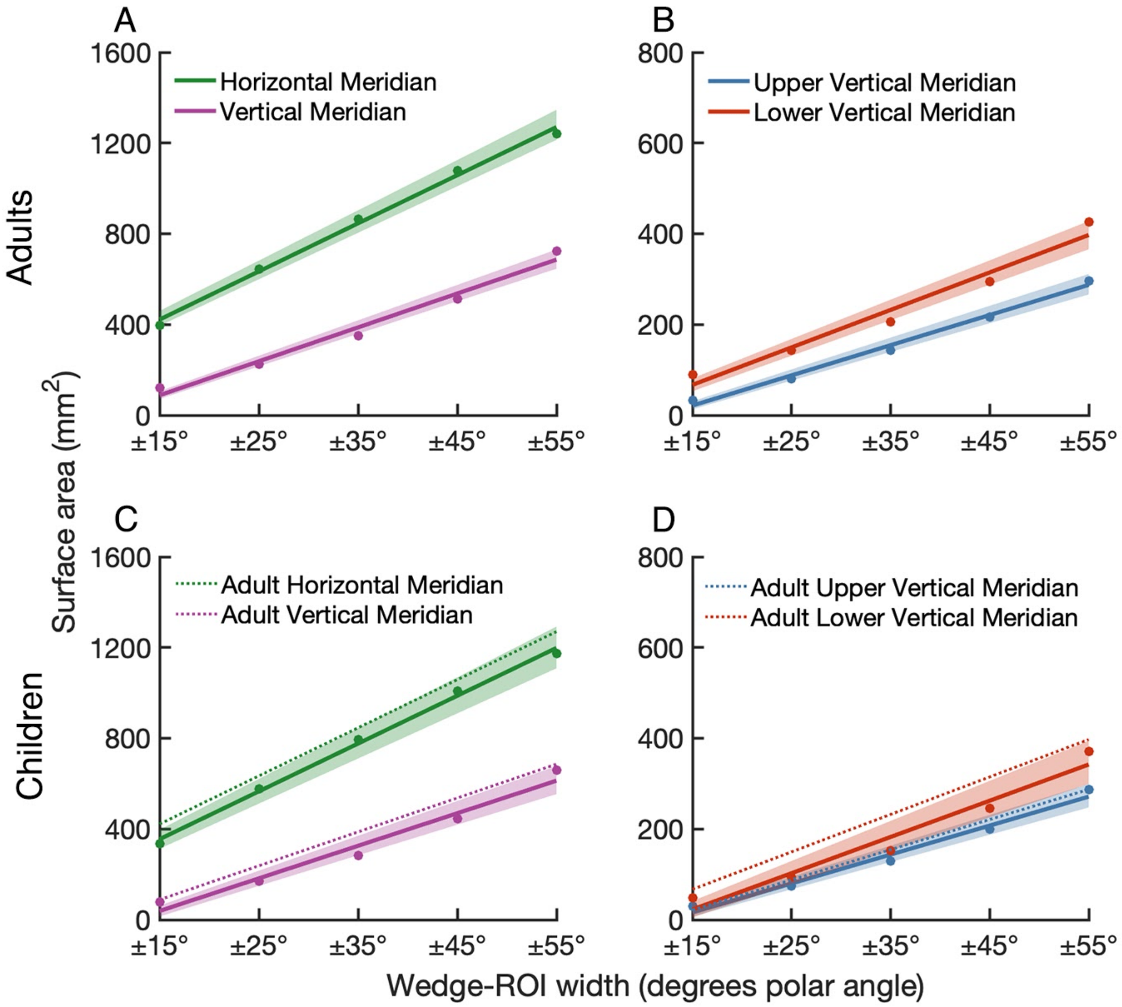


**SUPPLEMENTARY FIG. 6. Polar angle asymmetries in V1 surface area measured without enforcing contiguous regions and using raw pRF data.** Surface area measurements are calculated by identifying vertices with pRF centers between 1-7° eccentricity, and a range of increasing polar angle values. **(A and C)** V1 surface area ($mm^2$) measurements for vertices at a range of distances from the horizontal and vertical meridians adults (n=24) and children (n=25). **(B and D)** V1 surface area measurements for vertices at a range of distances from the upper and lower vertical meridians, for adults (n=24) and children (n=25). **(A-D)** Colored lines represent the average of 10,000 bootstrapped linear fits to the data. The colored data points represent the bootstrapped mean at each wedge-ROI width. The shaded error bar represents the 68% bootstrapped confidence interval of a linear fit to the data. The dotted lines in **(C)** and **(D)** are the adult surface area measurements replotted for comparison to children.

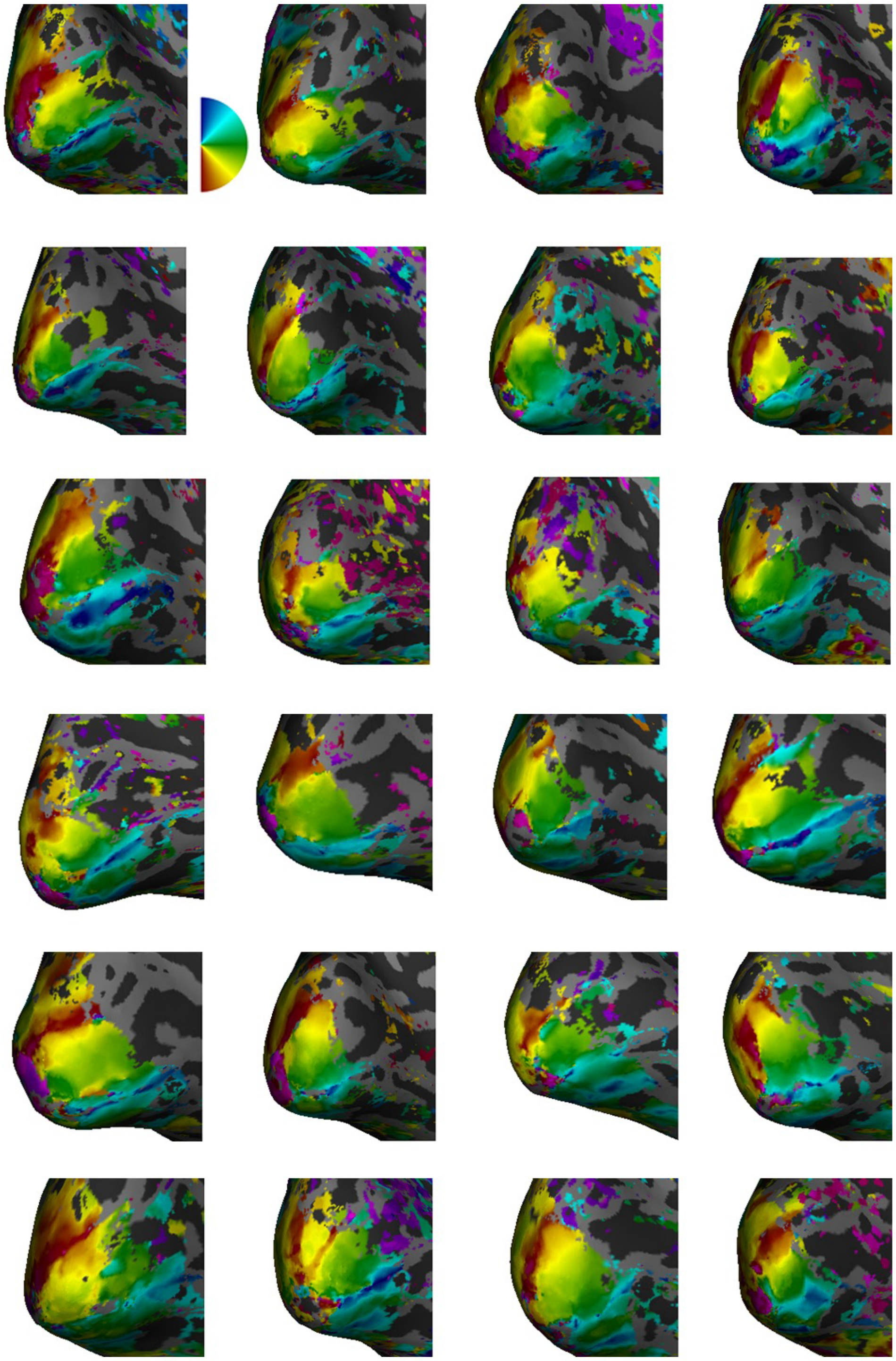

**SUPPLEMENTARY FIG. 7. Example left hemispheres from 24 adults.** Polar angle maps are projected onto the left hemispheres of inflated native surfaces that are angled to show V1. Red polar angle data represents the lower vertical meridian, blue data represents the upper vertical meridian, and green data represents the horizontal meridian.

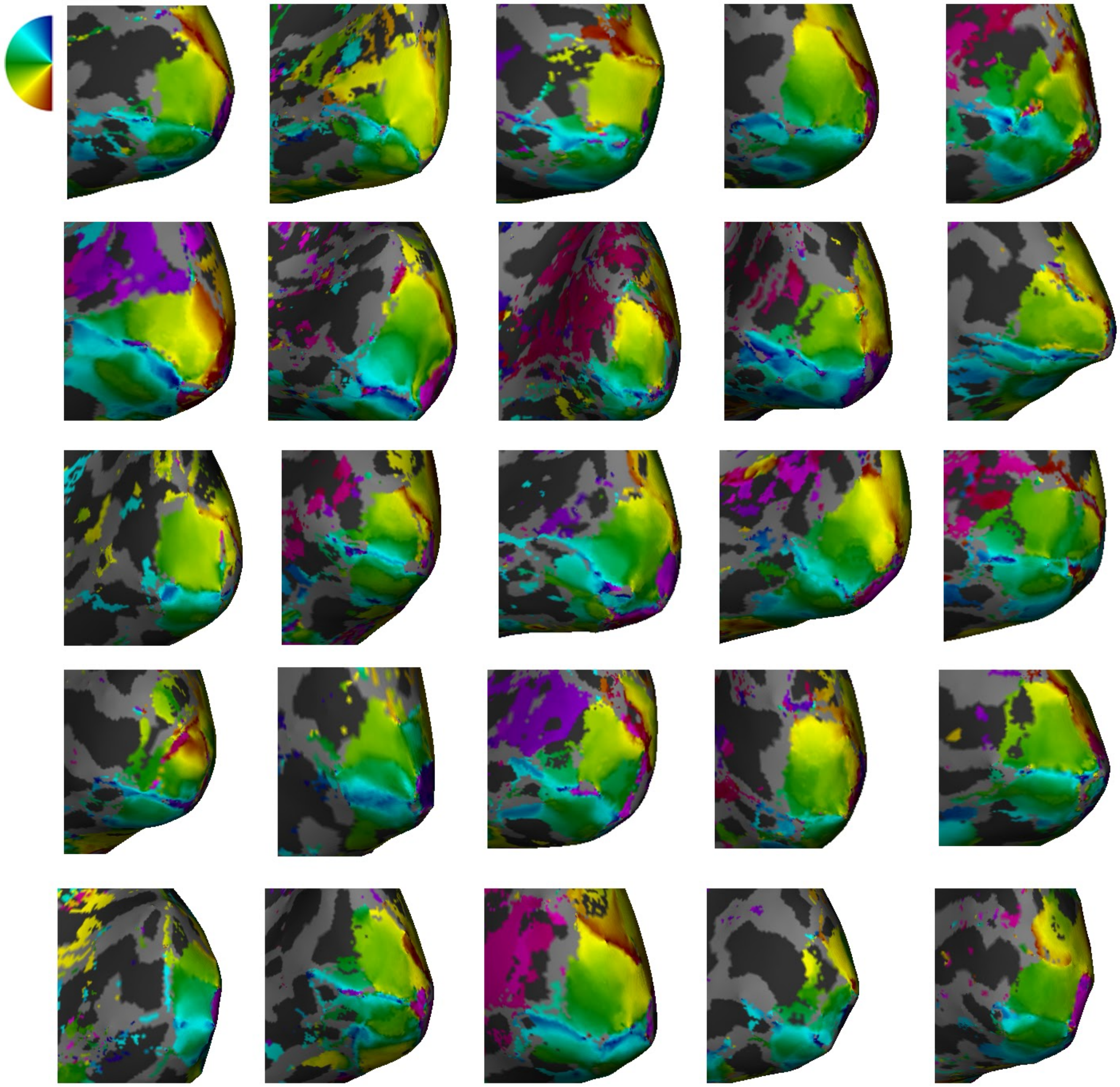

**SUPPLEMENTARY FIG. 8. Right hemisphere polar angle maps for all 25 children.** Polar angle maps projected onto the right hemisphere of the inflated native surfaces angled to show V1. Red polar angle data represents the lower vertical meridian, blue data represents the upper vertical meridian, and green data represents the horizontal meridian (see polar angle legend inset). Children had a thinner lower vertical meridian representation (red data) than adults, congruent with a reduced cortical representation of the lower vertical meridian of the visual field.

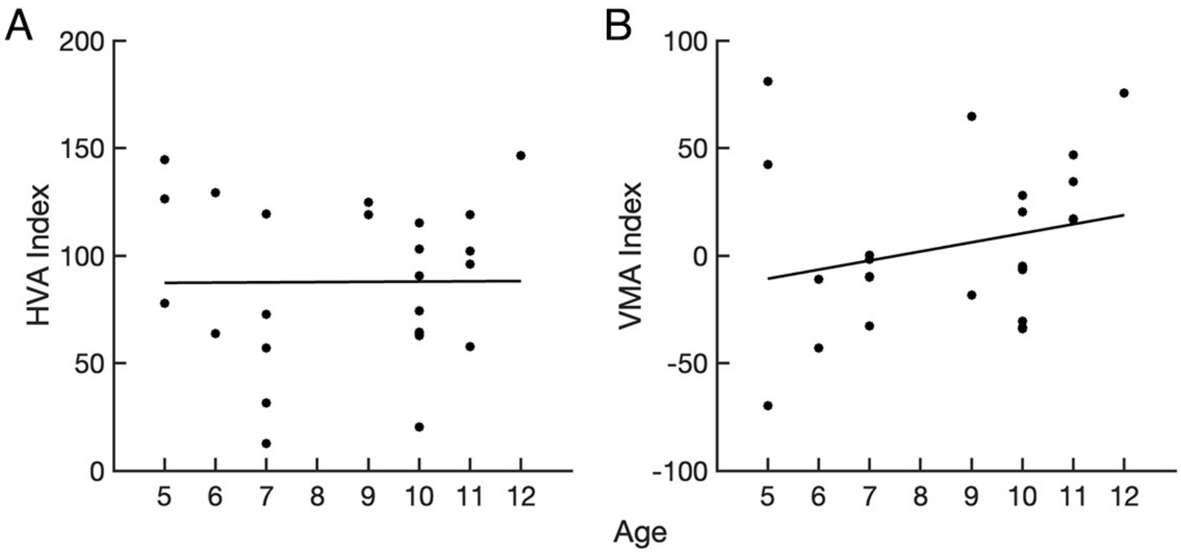


**SUPPLEMENTARY FIG. 9.** Spearman's correlations (two-tailed) of the magnitude of the **(A)** HVA index and **(B)** VMA index with age for children (n=25). The black line through the data is the line of best fit. Neither the magnitude of the HVA ($r$ = -0.0402 , $p$ = 0.848) or VMA ($r$=0.3068 , $p$=0.135) correlate with age for children.

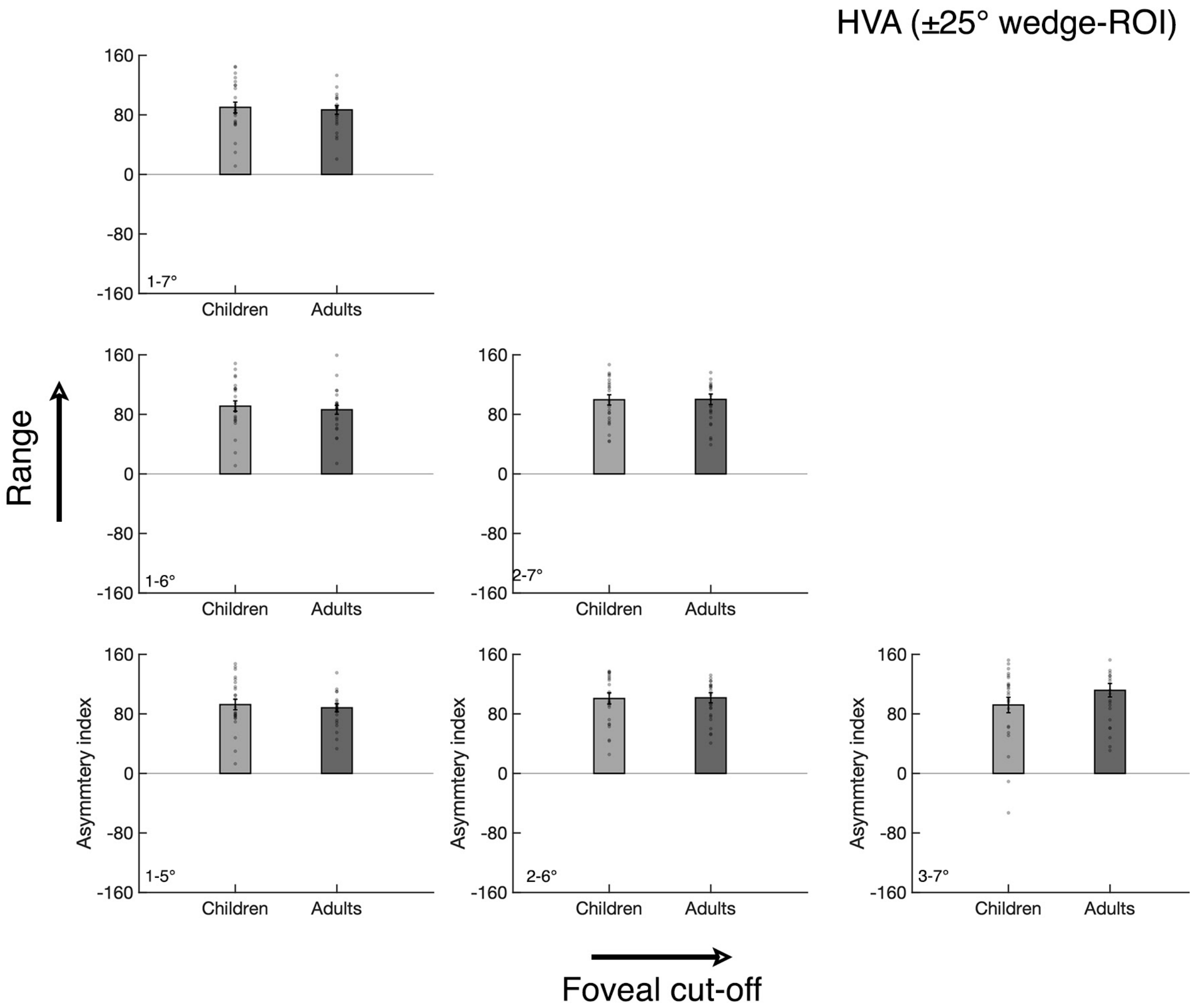


**SUPPLEMENTARY FIG. 10.** Cortical HVA measurements from different eccentricity ranges from the ±25° wedge-ROIs centered on the four polar angle meridians for children (n=25; light gray bars) and adults (n=24; dark gray bars). The data are restricted into varying eccentricity ranges. The 1-7° HVA is measured using the full wedge-ROI eccentricity range, as in the main text. Error bars indicate ±1 standard error of the mean (SEM).

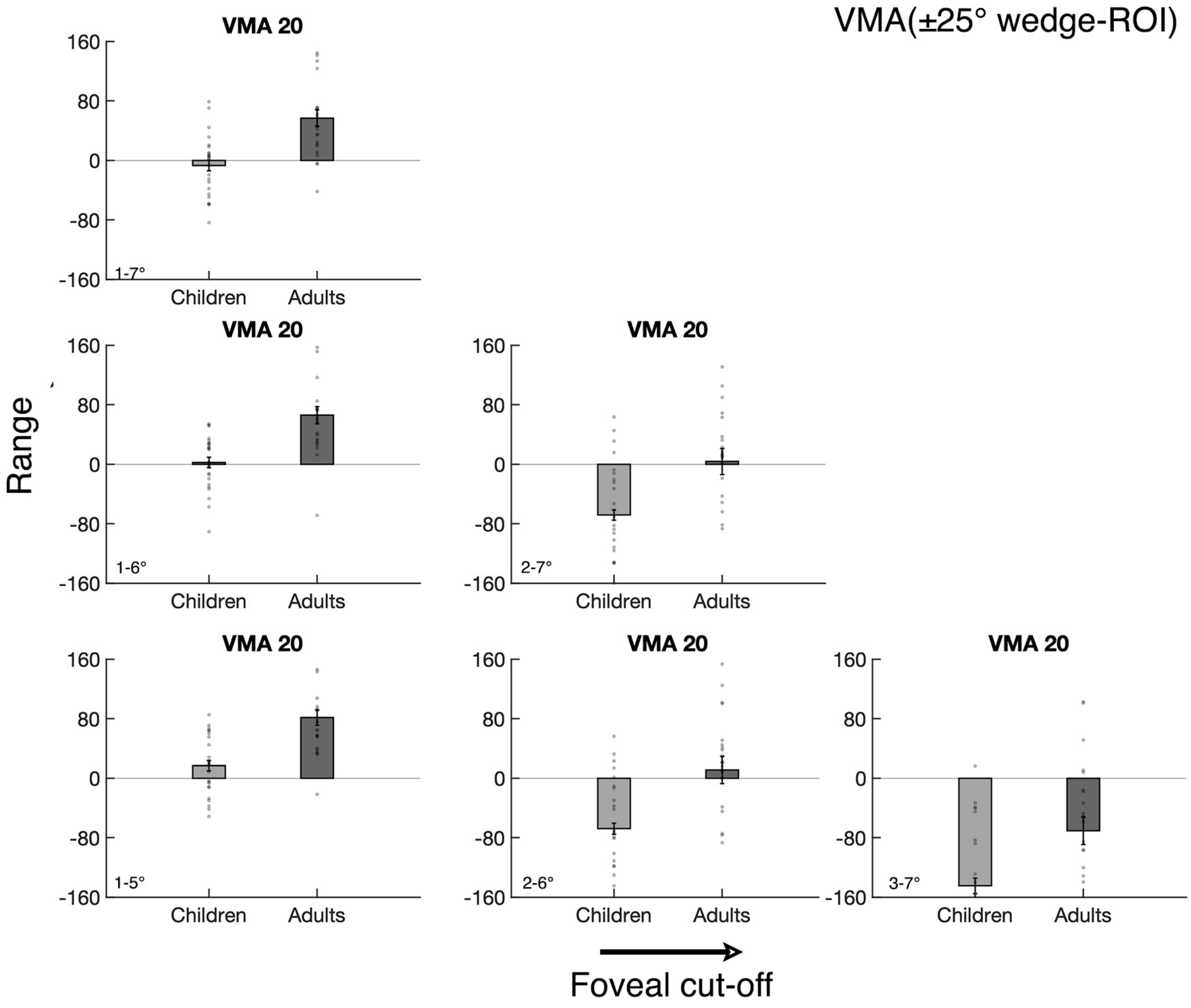


**SUPPLEMENTARY FIG. 11.** Cortical VMA measurements from different eccentricity ranges from the ±25° wedge-ROIs centered on the four polar angle meridians for children (n=25; light gray bars) and adults (n=24; dark gray bars). The data are restricted into varying eccentricity ranges. The 1-7° HVA is measured using the full wedge-ROI eccentricity range, as in the main text. Error bars indicate ±1 standard error of the mean (SEM).

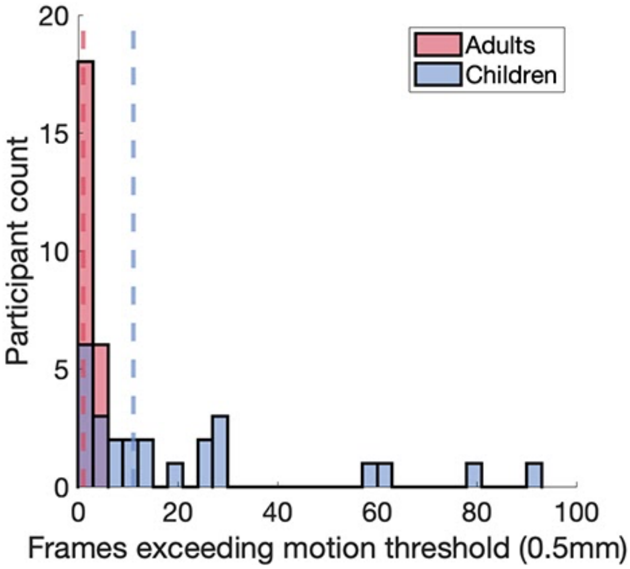


**SUPPLEMENTARY FIG. 12. Participant motion.** Within-scan motion presents the largest source of noise during fMRI[34] and children tend to move more than adults in the scanner[35,36]. To assess participant within-scan motion, we computed the number frames for which movement exceeded a motion threshold of 0.5mm. For each participant, this tally was summed across all four EPI scans (384 frames). The median tally for adults was 1 frame and the median tally for children was 10.5 frames. The plot shows histograms of the tally of frames in which motion exceeded threshold (0.5 mm) for adults (n=24, red) and children (n=25, blue). The dashed red and blue lines reflect the group median for adults and children, respectively.

**Supplementary Note 1:**

*Assessment of participant motion and the relationship between SNR and the cortical VMA*

We assessed whether differences in motion impact surface area measurements by correlating the surface area of each participant's V1, V2 and V3 ROIs with their within-scan motion. ROI surface area and participant motion did not correlate for adults (V1 $r$ = -0.11, V2 $r$ = 0.19 , V3 $r$ = 0.11, all $p > 0.1$) or children (V1 $r$ = 0.30 , V2 $r$ = 0.23 , V3 $r$ = 0.03 , all $p > 0.1$), indicating that the boundaries from polar angle and eccentricity maps are not impacted by participant motion. This is because the early visual field maps are defined by explicit polar angle and eccentricity boundaries, not by a signal-to-noise ratio threshold.

We also tested whether group differences in participant motion impact the magnitude of the HVA and VMA. Our goal here was to test whether the lack of a cortical VMA in children could be an artifact of within-scan motion. The magnitude of the cortical VMA did not correlate with participant motion, for adults ($r$ = -0.13, $p$ = 0.528) or children ($r$ = 0.07, $p$ = 0.730), indicating that the lack of cortical VMA in the children was not due to group differences in within-scan motion. Likewise, the magnitude of the cortical HVA did not correlate with participant motion for adults ($r$ = 0.09, $p$ = 0.680) or children ($r$ = -0.12, $p$ = 0.5462).

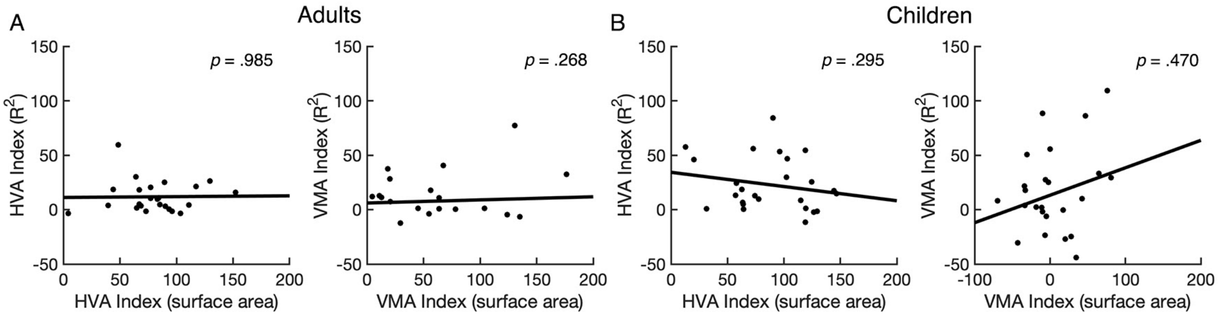


**SUPPLEMENTARY FIG 13.** Correlations between asymmetries in surface area and asymmetries in variance explained. Two-tailed Spearman's correlations show that the surface area HVA and $R^2$ HVA do not correlate in **(A)** adults (r = -0.01, p = 0.985) or **(B)** children (r = -0.22, p = 0.295). Likewise, the surface area VMA and $R^2$ VMA do not correlate in **(A)** adults (r = -0.23, p = 0.268) or **(B)** children (r = .15, p = 0.470).

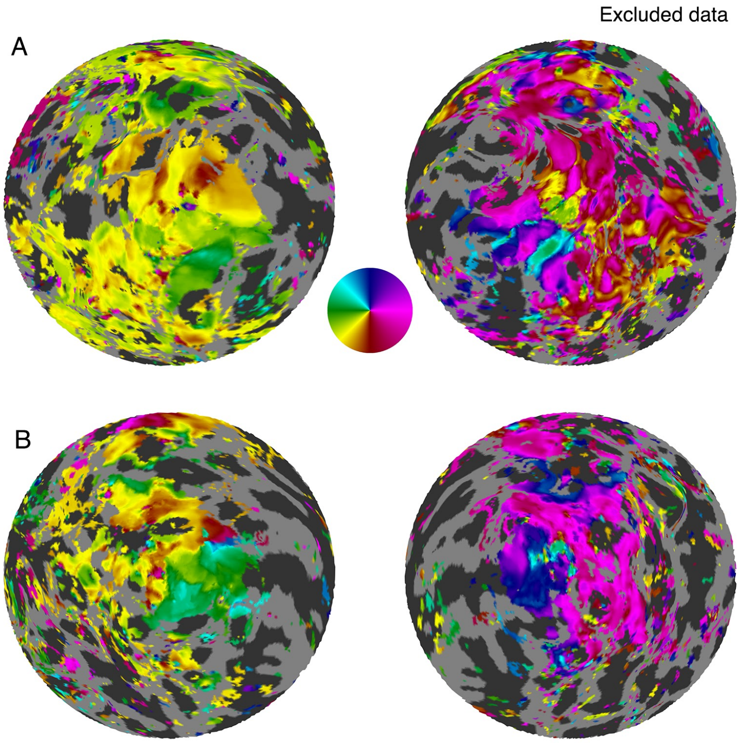


**SUPPLEMENTARY FIG 14.** Flatmaps of the polar angle pRF data from 2 excluded adults. The boundaries of V1 could not be reliably identified for these 2 participants, thus their data were not included in further analyses.
